# Nebulized 2-deoxylated glucose analogues inhibit respiratory viral infection in advanced *in vitro* airway models

**DOI:** 10.1101/2024.12.20.629606

**Authors:** Sarah K. Wideman, Laxmikant Wali, Vitalii Kovtunyk, Scharon Chou, Vanessa Gusel, Heta Telimaa, Chama Najmi, Delyana Stoeva, Johannes Stöckl, Guido A. Gualdoni, Anna-Dorothea Gorki, Snezana Radivojev

## Abstract

Respiratory viral infections caused by rhinoviruses (RVs), influenza A virus (IAV) and endemic corona viruses (HCoV) result in a serious strain on healthcare systems and public health, underscoring an urgent need for inhaled broad-spectrum antiviral therapies. However, their development is challenging, as no standardized *in vitro* methodologies that can fully replicate the *in vivo* environment have been established. In this work, we aimed to investigate the antiviral and anti-inflammatory effect of three 2-deoxylated glucose analogues (2-DGA): 2-deoxy-D-glucose, 2-fluoro-2-dexoy-D-glucose and 2-fluoro-2-dexoy-D-mannose (2-FDM), by utilizing advanced *in vitro* air-liquid interface (ALI) airway models. We demonstrated that commonly used ALI models have variable susceptibility to RV, IAV and HCoV infection. Further, we showed that 2-DGA have an anti-inflammatory effect and suppress respiratory viral replication in models mimicking the upper and lower respiratory airways. Moreover, we confirmed that 2-DGA can be delivered via nebulization *in vitro*, highlighting their potential to be used as broad-spectrum inhaled antivirals. Finally, our results demonstrate the importance of incorporating complex *in vitro* methodologies, such as primary cell ALI cultures and aerosol exposure, at an early stage of drug development.

## Introduction

Respiratory viral infections are a significant and growing global health challenge, particularly as a leading cause of morbidity and mortality among children and immunocompromised adults. These infections pose a considerable burden on healthcare systems and public health, as illustrated by the staggering statistic that on average, children have 6-8 and adults 2-4 colds per year^1^. In the USA alone, around 20-22 million days of absence from work and school are noted yearly^1^. Furthermore, viral infections are responsible for severe exacerbations in patients suffering from different chronic conditions such as asthma (50-80%), chronic obstructive pulmonary disease (COPD; 35%) and cystic fibrosis (15%)^2–4^.

Respiratory viruses encompass various families with distinct but overlapping effects on the respiratory tract. In this work, we focused on common and clinically relevant viruses: rhinovirus (RV), influenza A virus (IAV) and human coronavirus (HCoV). RVs are the most frequent cause of the common cold and primarily infect the upper respiratory tract^5^. Endemic HCoVs, typically cause mild to moderate upper respiratory tract infections^6^, while IAVs have a broader impact and often infect both the upper and lower respiratory tracts^7^. Their replication cycles have been well-studied, making them a good models for understanding viral infections^5,6,8–12^. Importantly, the pathology of these infections is significantly influenced by the host’s immune response, driving the severity of symptoms and exacerbations of pre-existing conditions^2,3,13^. Hence, understanding the fine balance between effective virus clearance and controlled immune activation is critical for managing and treating respiratory infections.

Inhaled antivirals offer several advantages over alternative delivery routes. By delivering the drug directly to the site of infection in the respiratory tract, antivirals can achieve higher local drug concentrations, while minimizing systemic exposure and side effects. This targeted delivery can lead to increased efficacy, faster symptom relief and reduced risk of complications. Additionally, inhalation bypasses the gastrointestinal tract and first-pass metabolism, improving bioavailability and efficacy compared to oral formulations. The non-invasive nature of inhaled therapies also makes them more patient-friendly compared to injected therapies, enhancing adherence and accessibility^14,15^. Still, the development of inhaled antivirals is not straightforward. It remains challenging to replicate key *in vivo* processes like drug deposition, clearance, dissolution, and permeability *in vitro*^16–18^. While simple monolayer cultures like Calu-3 or A549 cells provide basic insights^19,20^, advanced models like primary cell-based air liquid interface (ALI) cultures and organoids offer greater complexity, but they also incur higher costs and have increased variability^10,21^.

The primary aim of this study was to assess three 2-deoxylated glucose analogues (2-DGA): 2-deoxy-D-glucose (2-DG), 2-fluoro-2-dexoy-D-glucose (2-FDG) and 2-fluoro-2-dexoy-D-mannose (2-FDM), as inhaled antivirals against respiratory viruses. Like glucose, 2-DGA are taken up by cells and phosphorylated at the 6-carbon position by hexokinase. However, unlike glucose, 2-DGA cannot be further metabolised by glucose-6-phosphate isomerase and the phosphorylated 2-DGA accumulate within the cell and displace glucose ^22^. Viruses rely entirely on the host’s metabolism for their replication, and they are particularly dependent on glucose metabolism^23^. This dependency can be targeted by employing 2-DGA as host-targeted antiviral drugs^24–28^. The 2-DGA partially and reversibly inhibit glycolysis, and as a result, crucial glucose derivatives that are required for viral replication are depleted^25^. As non-infected host cells have lower baseline metabolism and are less reliant on glycolysis, 2-DGA are considered safe and non-toxic at moderate doses^29^.

Furthermore, we aimed to utilize advanced *in vitro* methodological approaches to increase translatability to the *in vivo* environment. These models were implemented with three specific objectives: (i) to demonstrate their ability to support infection with diverse respiratory viruses, (ii) to replicate the microenvironment of both the upper and lower respiratory tract, and (iii) to assess the importance of using more complex setups, such as aerosol exposure chambers, when designing therapies aimed at the lower respiratory tract.

## Results

### ALI airway models have variable susceptibility to respiratory viral infection

To evaluate the antiviral potential of 2-DGA, we aimed to use more complex and translationally relevant ALI cultures. In recent years, airway epithelial cells cultured at ALI have become prevalent in pulmonary drug development and respiratory infection studies. Although ALI cultures are considered a gold standard model, a variety of cell types are frequently used, and no single standardized model is agreed upon when studying respiratory viral infections^16,30–32^. Therefore, we systematically characterized the ALI culture infection kinetics of four common and clinically relevant virus strains, RV-A1B, RV-A16, IAV-H1N1 and HCoV-229E. Several widely used ALI models of the airway were infected, including both immortalized Calu-3 and HBEC3-KT cells and primary human nasal epithelial cells (HNEC) and human bronchial epithelial cells (HBEC).

Interestingly, not all ALI models were susceptible to all respiratory viruses. RV-A1B, a minor group RV that enters cells through the low-density lipoprotein receptor, infected neither HNEC nor HBEC (Fig. 1a, open symbols). However, RV-A1B replicated well in both immortalized cell models, i.e. Calu-3 and HBEC3-KT (Fig. 1a, closed symbols). In contrast, RV-A16, a major strain RV that uses ICAM-1 for cell entry, infected and replicated in all four ALI models (Fig. 1b). Most ALI models were also susceptible to HCoV-229E, except for HBEC3-KT cells (Fig. 1c). IAV-H1N1 was able to infect and replicate in all three tested models: Calu-3, HBEC3-KT and HBEC (Fig. 1d). IAV was not tested in HNEC as we were primarily interested in applying IAV as a model of lower respiratory tract infection. Notably, all four tested viruses were able to replicate in Calu-3 ALI cultures (Fig. 1).

**Figure 1.**
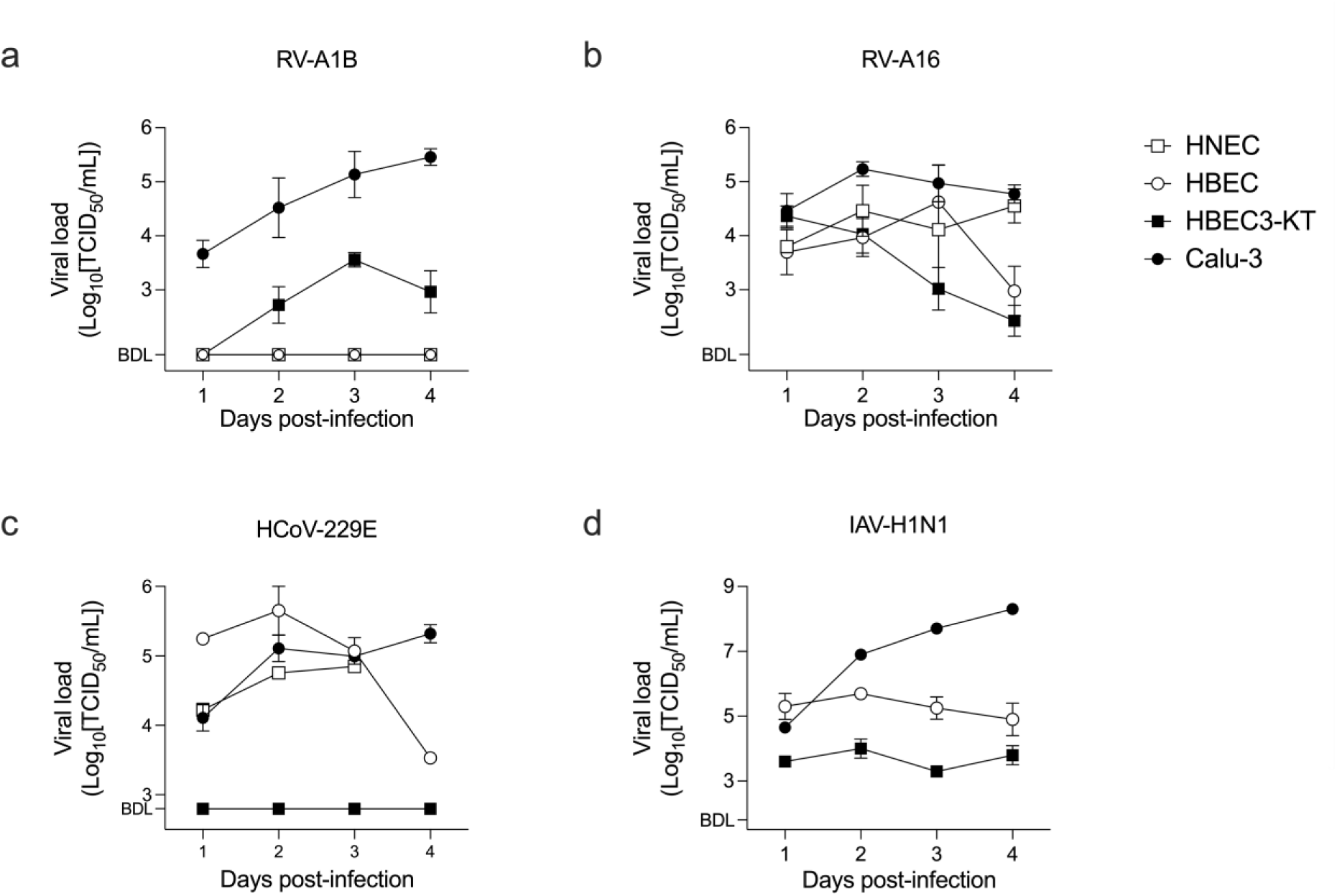
Human airway cells cultured at ALI respond differently to respiratory viral infection. Primary HNEC and HBEC, and immortalized HBEC3-KT and Calu-3 cells were infected with different respiratory viruses. Released infectious virus, i.e. the viral load, was measured over four days by titrating the TCID_50_ for (**a**) RV-A1B, (**b**) RV-A16, (**c**) HCoV-229E and (**d**) IAV subtype H1N1. Displaying mean, SEM, n=3-6, N=2 (a, b) or n=1-2, N=1 (c, d). **Abbreviations:** air-liquid-interface (ALI), human nasal epithelial cells (HNEC), human bronchial epithelial cells (HBEC), rhinovirus (RV), human coronavirus (HCoV), influenza A virus (IAV), below detection limit (BDL), median tissue culture infectious dose (TCID_50_).

Overall, the susceptibility to respiratory viral infection differs between airway ALI models, likely due to factors such as the different cell type composition, varying surface receptor expression and divergent innate antiviral responses. Based on the screening results, further assessments of the 2-DGA were performed in the context of RV-A16 infection using primary cells or Calu-3 cells.

### 2-DGA inhibit RV infection in primary cell models of upper and lower airway

We began by evaluating a 2-DG formulation intended for nasal spray use in the context of upper airway infection. Thus, ALI cultured HNEC were infected with RV-A16 and treated apically with a solution of 3.5% 2-DG or placebo (NaCl) over the course of two days (Fig. 2a). HNEC originating from two different donors were utilized to identify any potential donor-dependent effects.

**Figure 2.**
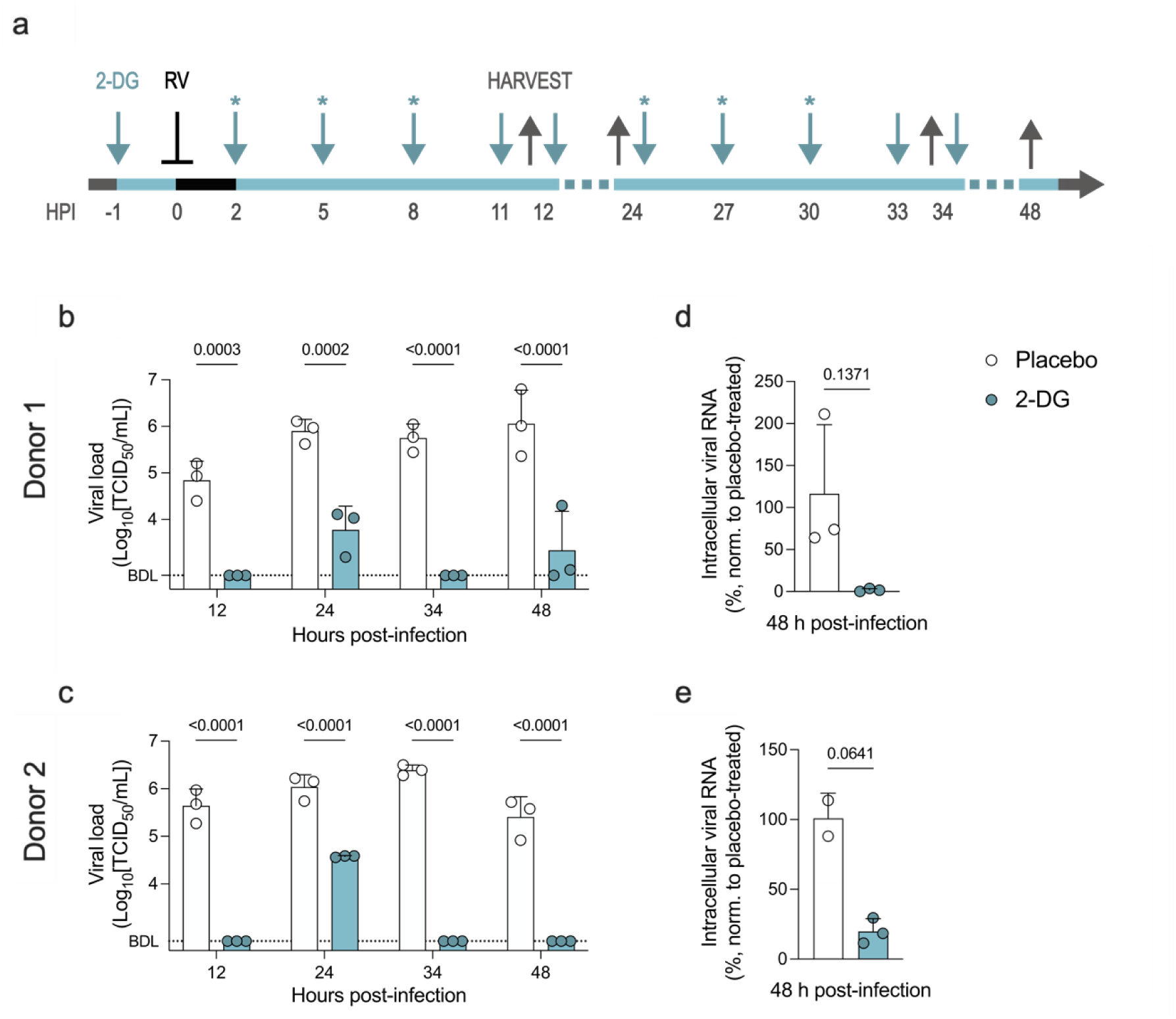
2-DG inhibits RV infection in HNEC from different donors. (**a**) ALI cultured HNEC were infected with mock or RV-A16 (⊥) and treated with a solution of 3.5% 2-DG (↓) at the indicated timepoints. The 2-DG treatment was removed 1 hour after application (*) or was left on the cells until samples were harvested (↑). (**b, c**) Released infectious virus, i.e. viral load. Displaying mean, SD, one-way ANOVA with Sidak’s test, n=3, representative of N=3. (**d, e**) Viral RNA levels relative to the placebo-treated control. Displaying mean, SD, Welch’s t-test, n=2-3, representative of N=3. **Abbreviations:** 2-deoxy-D-glucose (2-DG), air-liquid-interface (ALI), human nasal epithelial cells (HNEC), rhinovirus (RV), hours post-infection (HPI), below detection limit (BDL), median tissue culture infectious dose (TCID_50_).

2-DG treatment significantly reduced the viral load at all measured timepoints in both donors (Fig. 2b,c). Additionally, a statistically non-significant decrease in viral RNA was observed 48 hours after infection (Fig. 2d,e). The highest viral load in the 2-DG treated samples was observed 24 hours after infection, reasonably due to the long incubation time without treatment renewal. Importantly, a similar response to 2-DG treatment was observed in both donors, indicating that the antiviral effect of 2-DG is not donor dependent. Hence, 2-DG inhibits RV infection in a complex primary cell ALI model of the upper airway, further strengthening its potential application as an antiviral drug in a nasal spray formulation.

Next, we wanted to investigate the antiviral effect of all three metabolically inhibitory glucose analogues, 2-DG, 2-FDG and 2-FDM. Initial screening was performed in submerged immortalized human epithelial HeLa Ohio cells to assess if all three drugs had antiviral activity against RV. Thus, HeLa Ohio cells were infected with RV-B14 and treated with 2-DG, 2-FDG or 2-FDM for 7 hours. All three 2-DGA strongly inhibited viral replication to comparable levels (Supplementary Fig.1a).

To assess the antiviral effects of 2-DGA in lower airway cells, ALI cultured HBEC were infected with RV-A16 and treated apically with a solution of placebo (PBS) or 3.5% 2-DG, 2-FDG or 2-FDM for 27 hours (Fig. 3a). All three 2-DGA strongly suppressed viral replication (Fig. 3b) and decreased the release of infectious virus (Fig. 3c). The cell viability was high in all treatment groups, and neither infection nor drug treatment had a significant effect on viability (Fig. 3d). Hence, 2-DG, 2-FDG and 2-FDM have promising antiviral activity in primary lower airway cells and the possibility of utilizing them as inhaled antivirals should be further investigated.

**Figure 3.**
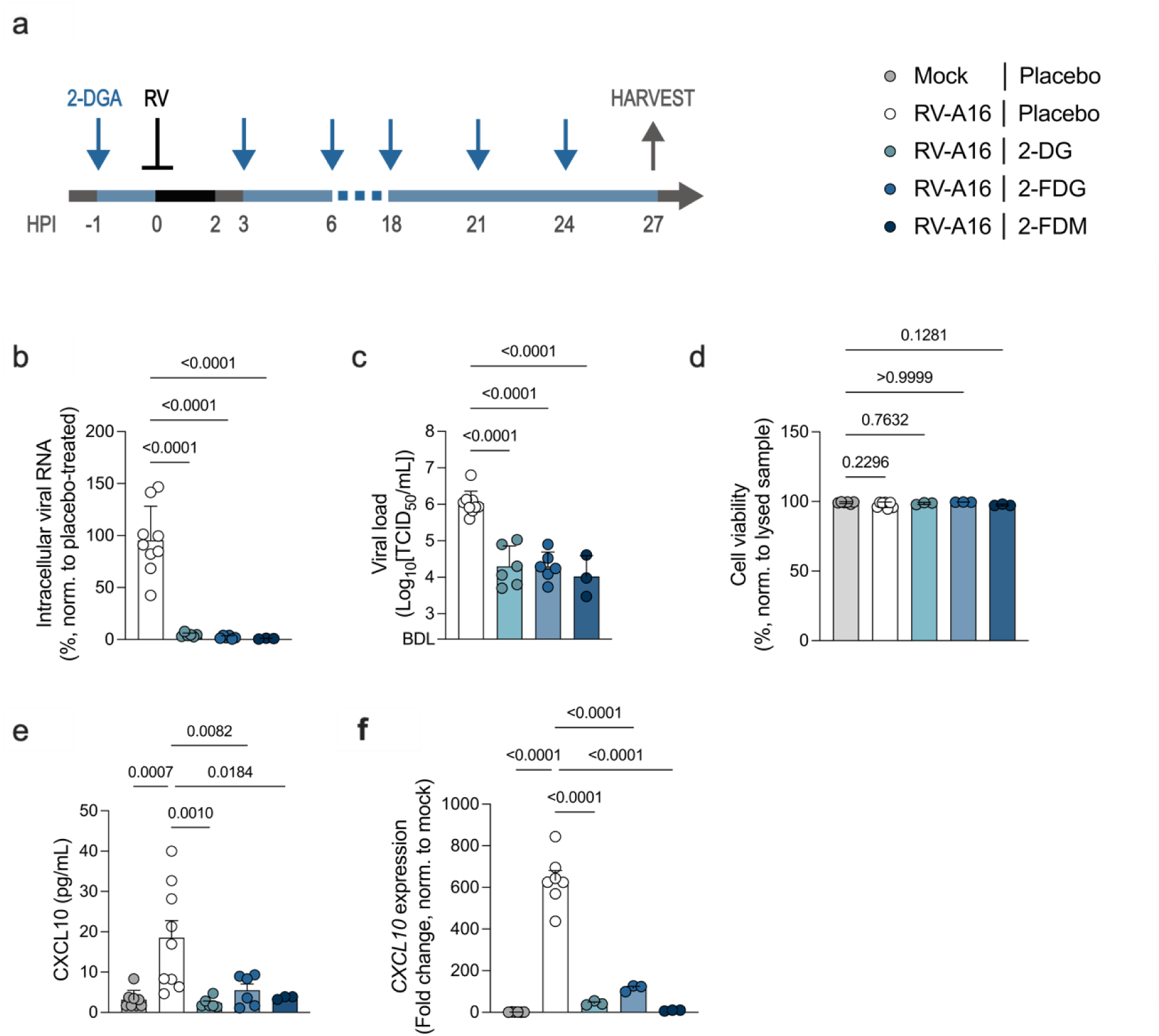
2-DGA have antiviral effects in RV-A16 infected HBEC. (**a**) ALI cultured HBEC were infected with mock or RV-A16 (⊥), treated with a solution of 3.5% 2-DGA (**↓**) and samples were harvested (↑) at the indicated timepoints. (**b**) Viral RNA relative to the placebo-treated controls. (**c**) Released infectious virus, i.e. viral load. (**d**) Cell viability measured via LDH release and normalised to a lysed HBEC sample. (**e**) CXCL10 levels in the basal medium. (**f**) *CXCL10* gene expression. Displaying mean, SD, one-way ANOVA with Tukey’s (b, c) or Dunnett’s (e, f) test or Kruskal-Wallis with Dunn’s test (d), n=3-9, N=3. **Abbreviations:** 2-deoxylated glucose analogues (2-DGA), 2-deoxy-D-glucose (2-DG), 2-fluoro-2-deoxy-D-glucose (2-FDG), 2-fluoro-2-deoxy-D-mannose (2-FDM), air-liquid-interface (ALI), human bronchial epithelial cells (HBEC), hours post-infection (HPI), lactate dehydrogenase (LDH), rhinovirus (RV), below detection limit (BDL), median tissue culture infectious dose (TCID_50_).

### 2-DGA have anti-inflammatory activity in HBEC and monocytes

Excess inflammation in response to infection is a key reason for symptoms and complications caused by respiratory viruses. Thus, the immunomodulatory activity of 2-DG, 2-FDG and 2-FDM was assessed by measuring the levels of pro-inflammatory cytokines and chemokines (CCL2, CCL5, IL-1β, IL-6, TNFα and CXCL10) secreted by RV-A16 infected HBEC. Out of the measured analytes, only CXCL10 was induced by RV-A16 infection in HBEC 27 hours after infection (Fig. 3e and Supplementary Fig. S2). However, 2-DGA treatment decreased CXCL10 to near baseline levels (Fig. 3e), indicating that 2-DGA treatment attenuates the pro-inflammatory response in RV-A16 infected HBEC. In addition, 2-DG, 2-FDG and 2-FDM treatment greatly repressed the transcription of *CXCL10* (Fig. 3f), further confirming our findings.

To determine if the 2-DGA have immunomodulatory properties independent of their antiviral effects, we assessed their effects on immortalized human monocytic THP-1 cells with a fluorescent NF-κB reporter gene^33^. The cells were activated with a synthetic viral RNA mimetic (R848) and treated with 2-DG, 2-FDG or 2-FDM. After 24 hours, NFκB translocation to the nucleus and cytokine production was measured. 2-DGA treatment significantly decreased the activation of key immune regulator and transcription factor NF-κB (Fig. 4a). Furthermore, treatment with 2-DG, 2-FDG and 2-FDM suppressed pro-inflammatory cytokine production by THP-1 cells (Fig. 4b). Therefore, 2-DG, 2-FDG and 2-FDM possess certain anti-inflammatory properties which may be beneficial in the treatment of respiratory viral infection.

**Figure 4.**
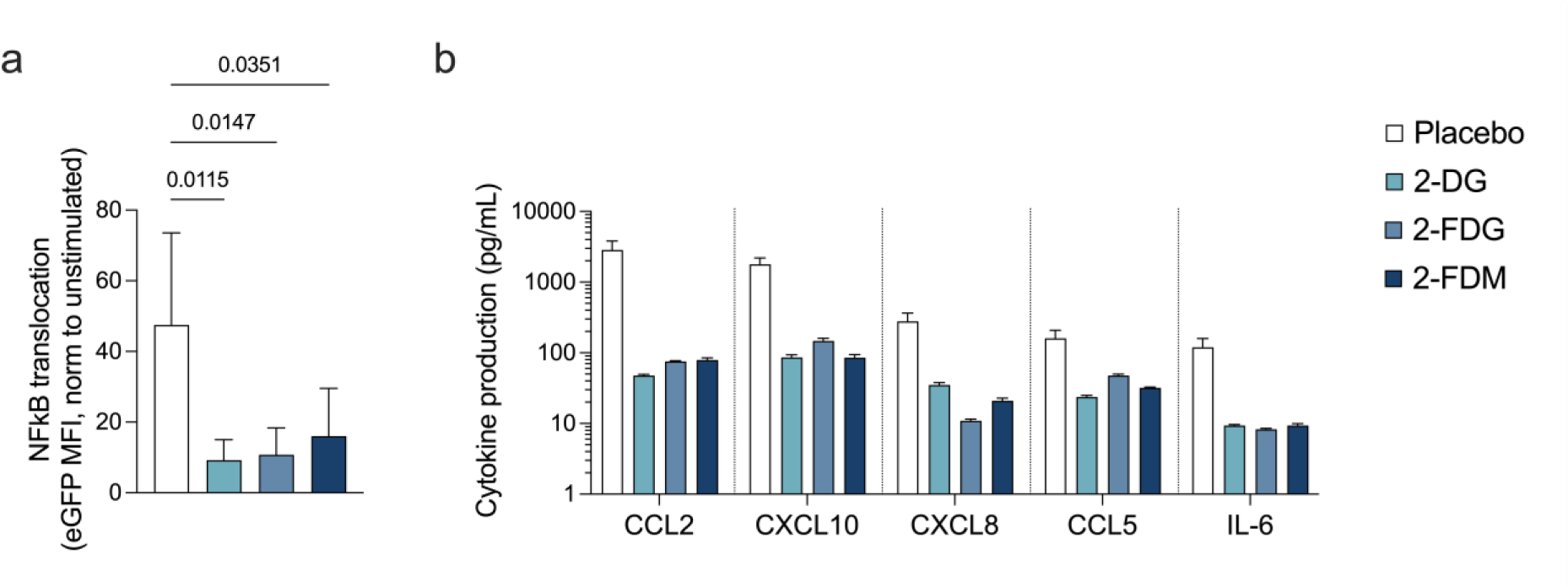
2-DGA have anti-inflammatory activity in human monocytic THP-1 cells. THP-1 cells were activated with R848 (30 µM) and treated with 2-DG, 2-FDG or 2-FDM (30 mM) for 24 hours. (**a**) NFκB translocation. The data was normalised to a control that was not treated with R848 (unstimulated). Displaying mean, SD, one-way ANOVA with Dunnett’s test, n=4, N=2. (**b**) Pro-inflammatory cytokine levels in THP-1 culture medium. Displaying mean and range, n=2, N=1. **Abbreviations:** 2-deoxylated glucose analogues (2-DGA), 2-deoxy-D-glucose (2-DG), 2-fluoro-2-deoxy-D-glucose (2-FDG) and 2-fluoro-2-deoxy-D-mannose (2-FDM), enhanced green fluorescent protein (eGFP), median fluorescence intensity (MFI).

### 2-FDG can be efficiently delivered through nebulization *in vitro*

Nebulizers are one type of inhalation device that can be used to deliver drugs to the lower airways. To replicate nebulizer mediated drug delivery in the pre-clinical assessment of the 2-DGA, vibrating mesh nebulizers and an *in vitro* aerosol exposure chamber were employed.

Firstly, we performed experiments to compare the drug uptake characteristics of 2-FDG delivered as aerosols, following nebulization, or delivered as a solution pipetted directly onto the cells. Nebulized 2-FDG was internalised by Calu-3 cells quickly, compared to a solution of 2-FDG, as higher levels of intracellular 2-FDG-6-phosphate (2-FDG-6P) were detected in cells treated with nebulized 2-FDG after one hour (Fig. 5a). However, after two hours the cellular uptake of 2-FDG was comparable, regardless of the delivery method (Fig. 5a). Although aerosol delivery of 2-FDG does not result in higher peak intracellular levels, it is faster and therefore more efficient. This may be particularly relevant for antivirals against respiratory infections, as they need to act quickly to prevent or slow down virus replication.

**Figure 5.**
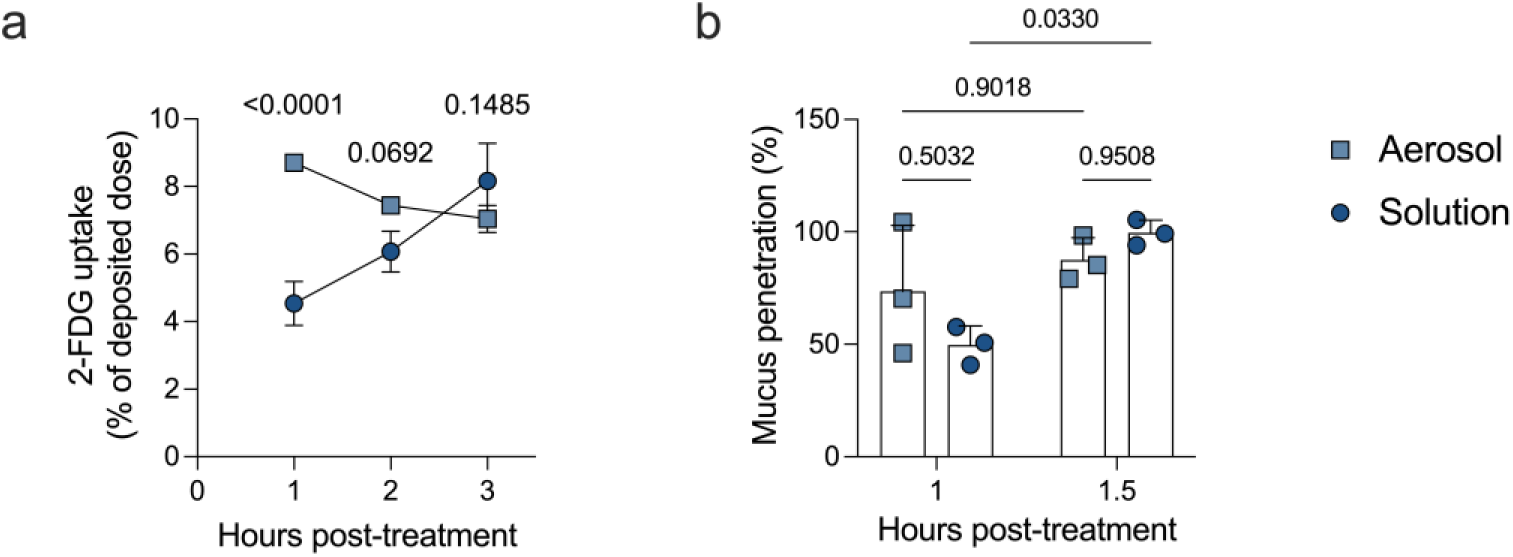
2-FDG delivered as aerosols has faster uptake kinetics than 2-FDG delivered as a solution. (**a**) Calu-3 cells cultured at ALI were treated with nebulized 2-FDG or a solution of 2-FDG. Cellular 2-FDG uptake, presented as the proportion of deposited 2-FDG that was internalised at each timepoint. Displaying mean, SD, one-way ANOVA with Sidak’s test (comparing Aerosol and Solution), n=3, representative of N=2. (**b)** Mucus was applied to empty transwells that were subsequently exposed to 2-FDG aerosols or a 2-FDG solution. 2-FDG mucus penetration relative to the delivered dose. Displaying mean, SD, one-way ANOVA with Sidak’s test (comparing all groups), n=3, representative of N=2. **Abbreviations:** 2-fluoro-2-deoxy-D-glucose (2-FDG), air-liquid-interface (ALI).

Drugs that are delivered locally to the respiratory tract need to pass through the mucus layer that covers and protects the epithelium within the conducting airways. Hence, we further investigated the mucus penetration kinetics of 2-FDG delivered via nebulization or as a solution. A cell-free system was utilized to understand drug-mucus interactions independent from varying cellular drug uptake efficiencies. Mucus was added to empty transwells that were subsequently exposed to 2-FDG delivered as aerosols or a solution. No statistically significant difference between aerosol and solution mucus penetration after 1 or 1.5 hours was observed (Fig. 5b). However, when delivered as a solution, more 2-FDG had penetrated the mucus layer after 1.5 hours than after 1 hour. In contrast, there was no difference in the mucus penetration of nebulized 2-FDG at the two timepoints. Hence, it is possible that a difference in mucous penetration may have been observed at earlier timepoints, however, due to technical constraints related to the aerosol exposure chamber this could not be assessed.

Taken together, we have shown that 2-FDG can be efficiently delivered *in vitro* through nebulization. Furthermore, there are subtle differences in the drug uptake characteristics between 2-FDG delivered as aerosols and as a solution.

### Nebulized 2-FDG inhibits respiratory viral infection

Having shown that 2-FDG can be delivered through nebulization *in vitro*, we next evaluated the antiviral effect of nebulized 2-FDG. Solution 2-FDG treatment was also included to understand whether the *in vitro* delivery method affects the antiviral efficacy. Two respiratory viruses were utilized to assess if nebulized 2-FDG could have a broad-spectrum antiviral effect.

Hence, Calu-3 cells cultured at ALI were infected with RV-A16 or HCoV-229E and treated with placebo (PBS) or 2-FDG, delivered as aerosols or as a solution, over the course of ten hours (Fig. 6a). 2-FDG inhibited early RV-A16 replication without causing any cytotoxic effects, independent of the delivery method (Fig. 6b,c). Moreover, the barrier integrity, a primary function of respiratory epithelium, was maintained in all treatment groups following infection and aerosol drug treatment (Fig. 5d). In addition, nebulized 2-DG supressed RV-A16 replication ten hours after infection (Supplementary Fig. S3). HCoV-229E replication was inhibited by both nebulized and solution 2-FDG treatment (Fig. 6e). High cell viability was maintained in all treatment groups upon HCoV-229E infection and drug treatment (Fig. 6f). Therefore, nebulized 2-FDG has broad-spectrum antiviral activity against two biologically distinct respiratory viruses.

**Figure 6.**
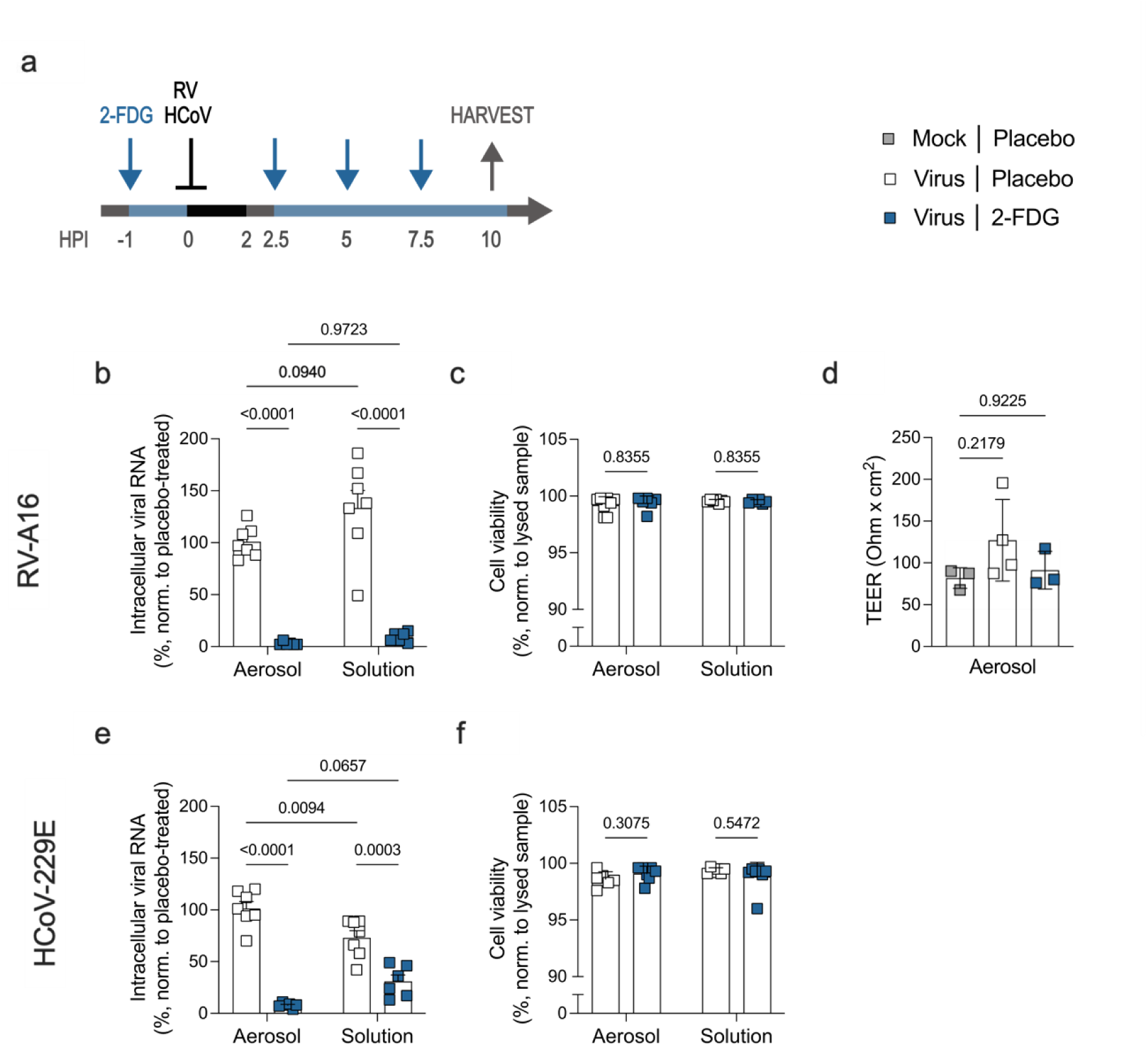
Nebulized 2-FDG inhibits RV and HCoV replication. (**a**) Calu-3 cells cultured at ALI were infected with mock, RV-A16 or HCoV-229E (⊥) and treated with nebulized 3.5% 2-FDG or an equivalent dose of 2-FDG delivered as a solution (**↓**), at the indicated timepoints. Samples were harvested (↑) 10 HPI. (**b**) RV-A16 viral RNA relative to the placebo-treated control. Displaying mean, SD, two-way ANOVA with Tukey’s test (comparing all groups), n=6-7, N=2. (**c**) Cell viability measured via LDH release and normalised to a lysed Calu-3 sample. Displaying mean, SD, multiple Mann-Whitney tests with Holm-Sidak’s test, n=6-7, N=2. (**d**) Barrier integrity as determined by measuring the TEER. Displaying mean, SD, one-way ANOVA with Dunnett’s test (comparing all groups), n=3-4, N=1. (**e**) HCoV-229E viral RNA relative to the placebo-treated control. Displaying mean, SD, two-way ANOVA with Tukey’s test (comparing all groups), n=6-7, N=2. (**f**) Cell viability measured via LDH release and normalised to a lysed Calu-3 sample. Displaying mean, SD, multiple Mann-Whitney tests with Holm-Sidak’s test, n=6-7, N=2. **Abbreviations:** 2-fluoro-2-deoxy-D-glucose (2-FDG), air-liquid-interface (ALI), human coronavirus (HCoV), hours post-infection (HPI), lactate dehydrogenase (LDH), rhinovirus (RV), trans-epithelial electrical resistance (TEER).

Interestingly, the treatment delivery method affected viral replication to some degree. For both RV-A16 and HCoV-229E infected cells, viral replication was different in the aerosol and solution placebo-treated groups. However, nebulized placebo treatment had opposing effects on RV-A16 and HCoV-229E, with a trend toward lower RV-A16 replication and significantly higher HCoV-229E replication (Fig. 6b,e). These observations underline the importance of using translatable models that replicate the intended delivery method for pre-clinical drug development studies.

### Nebulized 2-DGA inhibits RV infection in primary HBEC

We ultimately combined the primary HBEC ALI model with nebulizer mediated drug delivery. To this end, HBEC were infected with RV-A16 and treated with nebulized placebo (PBS), 2-DG or 2-FDG for 9 hours (Fig. 7a).

**Figure 7.**
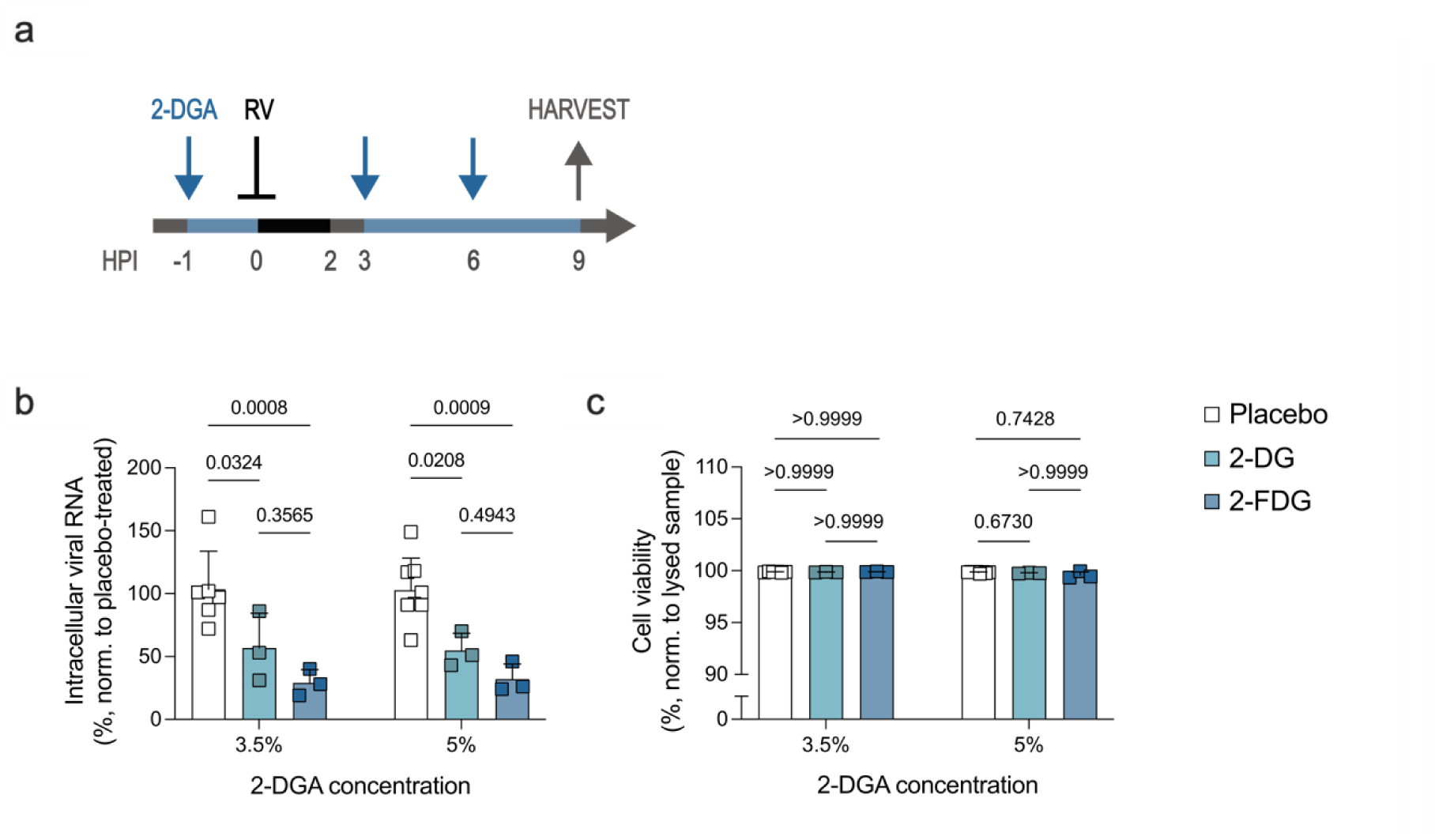
Nebulized 2-DGA inhibit RV replication in HBEC. (**a**) HBEC cultured at ALI were infected with RV-A16 (⊥), treated with nebulized 2-DG or 2-FDG (**↓**), and samples were harvested (↑) at the indicated timepoints. (**b**) RV-A16 viral RNA relative to the placebo-treated control. Displaying mean, SD, two-way ANOVA with Tukey’s test, n=3-8, N=4. (**c**) Cell viability measured via LDH release and normalised to a lysed HBEC sample. Displaying mean, SD, Kruskal-Wallis test with Dunn’s test, n=3-8, N=4. Statistical testing was performed separately for the two different drug concentrations. **Abbreviations:** 2-deoxylated glucose analogues (2-DGA), 2-deoxy-D-glucose (2-DG), 2-fluoro-2-deoxy-D-glucose (2-FDG), air-liquid-interface (ALI), human bronchial epithelial cells (HBEC), hours post-infection (HPI), lactate dehydrogenase (LDH), rhinovirus (RV).

Treatment with nebulized 3.5% 2-DG and 2-FDG significantly decreased RV-A16 replication (Fig. 7b). However, there was no significant difference in the antiviral efficacy of 3.5% 2-DG and 2-FDG (Fig. 7b). As the antiviral effect of 3.5% 2-FDG was less pronounced in HBEC than in Calu-3 cells (Fig. 6b & Fig. 7b), we next assessed if increasing the treatment dose would improve the antiviral effect. Thus, RV-A16 infected HBEC were treated with nebulized 5% 2-DG and 2-FDG (Fig. 7a). A comparable antiviral effect was achieved when treating with 5% as with 3.5% nebulized 2-DG and 2-FDG (Fig. 7b), indicating that moderate increases in treatment dose do not impact the antiviral efficacy. Cell viability remained high in all treatment groups regardless of the treatment dose (Fig. 7c). Considering the osmolality limitations of inhaled nebulized therapies and the lack of improved antiviral efficacy of 5% compared to 3.5% 2-DG and 2-FDG, more concentrated 2-DGA solutions were not evaluated.

In summary, 2-DG and 2-FDG are promising antiviral compounds that have antiviral activity against respiratory viruses in a range of *in vitro* models. Nebulizer delivery of 2-DGA appears to be effective, although formulation optimization may be required to ensure sufficient doses are delivered to cells within the complex lung environment.

## Discussion

Respiratory viral infections have a devastating impact on public health and novel effective broad-spectrum antivirals are greatly needed. However, antiviral development is complex and faced with numerous challenges, including the current state of available *in vitro* methodologies. Here, we utilized advanced primary cell-based ALI models of the human airway in combination with an aerosol exposure chamber to evaluate 2-DGA as inhaled antivirals. We have shown that 2-DGA can be delivered via nebulization *in vitro* and that they are promising antiviral drug candidates with anti-inflammatory properties and antiviral activity against common respiratory viruses.

We chose to assess the antiviral activity of 2-DGA in ALI cultures because they more closely mimic the human airway, natural viral infection and inhaled drug delivery, compared to submerged cell cultures^34^. Regardless, we found that several cell types that are commonly used for ALI cultures did not support infection with all tested respiratory viruses. Investigating the underlying cause was beyond the scope of this study. However, inherent host factors, such as varying viral entry receptor expression and innate antiviral immune responses, were likely at play. Additional complexity is introduced in the case of primary cells, where susceptibility to viral infection can also be donor specific^35^. This may explain why we observed no RV-A1B replication in HNEC and HBEC, even though RV-A1 infection of HBEC has previously been reported^36^. Taken together, these findings highlight that no model can fully replicate all features of the human airway, which should be considered carefully when choosing an experimental model.

The antiviral properties of 2-DG have been studied in cells cultured under standard submerged conditions since the 1950s^28^, including its activity against respiratory viruses^24,25,37,38^. However, until now 2-FDG and 2-FDM had not been investigated as antivirals against respiratory infections. We have shown, for the first time, that 2-DG, 2-FDG and 2-FDM inhibit viral replication in RV infected ALI cultured primary HBEC, without inducing any cytotoxic effects.

Hence, this study further establishes 2-DGA as a class of compounds that possess antiviral activity against respiratory viruses, thereby opening the door for further development and optimization. Importantly, 2-FDG inhibited the replication of both RV and HCoV, even though they belong to different families and have important biological differences, such as their entry receptors, virion size and whether they are covered by a capsid or an envelope. Coupled with previous findings^24,28,37–39^, this demonstrates that 2-DGA have the potential to be utilized as broad-spectrum antivirals against respiratory viruses.

Furthermore, the anti-inflammatory effects observed in infected HBEC and THP-1 cells treated with 2-DG, 2-FDG or 2-FDM indicate that the benefits of 2-DGA may go beyond antiviral. This is in line with the current understanding that pro-inflammatory immune responses are highly glycolytic processes^40^. Dysregulated immune responses to a respiratory viral infections can have severe repercussions, causing acute respiratory distress syndrome or lung disease exacerbations^2,3,13^. Hence, antivirals with additional anti-inflammatory effects could potentially prevent or ameliorate such severe symptoms.

Overall, the use of an aerosol exposure chamber to model *in vivo* nebulizer mediated drug delivery *in vitro* was a critical aspect of this study. We found that the uptake kinetics of nebulized 2-FDG were faster, as previously established for anti-cancer medication Bortezomib^41^. Nevertheless, the antiviral efficacy of aerosol and solution 2-FDG treatment was equivalent in RV and HCoV infected Calu-3 cells. Hence, utilizing the same *in vitro* method as the intended *in vivo* method for drug delivery appears to be important for detecting subtle differences, which can be consequential when defining the desired therapeutic approach.

We observed that nebulized 2-FDG treatment was less effective in RV infected HBEC than in Calu-3 cells. Calu-3 cells are highly glycolytic cancer derived cells^42^, meaning that they are more likely to be strongly affected by glycolytic inhibitors. Moreover, Calu-3 cells form a monolayer of cells when cultured at ALI, in contrast to HBEC that form a notably thicker cell layer with differentiated cells. HBEC also secrete thick mucus that could interact with the 2-DGA. It is known that the presence of mucus can decrease or completely abolish drug permeability and uptake, putatively due to drug-mucus interactions^18,43^. Ultimately, HBEC reflect the *in vivo* environment more authentically and the restricted antiviral effect of 2-FDG underlines the challenges inherent to delivering inhaled drugs at sufficient quantities to the lung. Since a strong antiviral effect can be achieved in HBEC, as seen with solution treatment, formulation optimization or alternative delivery approaches may be required to achieve the desired antiviral effect of inhaled 2-DGA *in vivo*.

Herein, we aimed to evaluate the antiviral activity of nebulized 2-DGA against respiratory viral infections using advanced *in vitro* models of the airway. We have found that advanced models such primary cell ALI cultures and an aerosol exposure chamber can be successfully implemented to perform pre-clinical assessments of antiviral drugs. Moreover, 2-DGA are promising drug candidates with anti-inflammatory and broad-spectrum antiviral properties.

## Methods

Detailed information on the origin of cells, viruses and key materials and equipment (Supplementary Table S1), as well as cell culture medium compositions (Supplementary Table S2) can be found in the Supplementary Information.

### Statement of ethical approval

According to Austrian law and the ’World Medical Association Declaration of Helsinki - Ethical Principles for Medical Research Involving Human Subjects’, the use of anonymised, non-identifiable cells obtained from a commercial provider do not require specific ethical approval. Primary human cells were purchased from Epithelix and were originally obtained with informed consent as part of processes that had ethical review and approval.

### Submerged cell culture

HeLa Ohio, MRC-5, MDCK and THP-1 NF-κB eGFP reporter cells were cultured in tissue culture-treated flasks at 37°C, 5% CO_2_, 85% humidity using cell-specific maintenance medium (see Supplementary Table S2). Adherent cells were passaged or used for experiments upon reaching 70-90% confluency and THP-1 cells were passaged 1-2 times per week.

### Air-liquid interface cultures

HNEC, HBEC, HBEC3-KT and Calu-3 cells were cultured at 37°C, 5% CO_2_, 85% humidity using cell-specific and differentiation-specific culture medium.

For submerged expansion culture in tissue culture-treated flasks, HNEC and HBEC were cultured in PneumaCult-Ex Plus medium, Calu-3 cells in Calu-3 medium and HBEC3-KT cells were grown in gelatine (0.1%) coated flasks using Keratinocyte-SFM medium. The cells were maintained until 70-90% confluency was reached, before detaching and seeding into 24-well plate transwell inserts. HNEC and HBEC were detached using an animal component-free dissociation kit, and HBEC3-KT and Calu-3 were detached using Accutase. Cells were seeded at a density of 6.0×10^4^ – 8.0×10^4^ cells per transwell insert. HNEC, HBEC and Calu-3 cells were maintained in the same medium as previously, while HBEC3-KT cells were transferred into in PneumaCult-Ex Plus medium.

At confluence, the cultures were airlifted by removing medium from the apical compartment and supplying cell-specific ALI medium to the basal compartment. HNEC, HBEC and HBEC3-KT cells were cultured in PneumaCult-ALI medium and Calu-3 cells in Calu-3 medium. The medium was refreshed every 2-4 days. Mucus was removed from HNEC, HBEC, and HBEC3-KT cultures through weekly apical washes with DPBS. Apical secretions were removed from Calu-3 ALI cultures every 3-4 days by aspiration.

### Viral infection & 2-DGA treatment

All viral infections and drug treatments were performed at 34°C, 5% CO_2_, and 85% humidity.

To infect HeLa Ohio cells with RV-B14, the cells were plated at a density of 1.5×10^5^ cells/well in a 24-well plate in 500 µL HeLa Ohio experiment medium. The following day, the cells were washed with PBS and the medium was replaced before infecting with 1.5×10^3^ TCID_50_/well of RV-B14. Simultaneously, treatment was initiated with 10 mM 2-DG, 2-FDG or 2-FDM diluted in culture medium, or with placebo (medium only). The cells were incubated for 7 h before washing with medium and collecting cell lysates.

ALI cultures were kept in their respective maintenance medium, except for HBEC, which were transferred into HBEC infection medium. ALI cultures were infected with 1×10^4^ TCID_50_/well of RV-A1B, 1-5×10^4^ TCID_50_/well of RV-A16, 0.75×10^4^ TCID_50_/well of HCoV-229E or 1.5×10^4^ TCID_50_/well of IAV-H1N1. For RV-A16, 1×10^4^ TCID_50_/well was used in conventional solution treatment experiments, while 5×10^4^ TCID_50_/well was used for nebulized treatment experiments. Infections were performed by adding 40 µL virus diluted in PBS, or PBS alone for mock infected controls, to the apical compartment of each ALI culture and incubating for 2 h. Afterwards, the cultures were washed six times with DPBS to remove non-internalised virus. Apical washes with DPBS were performed to collect samples for TCID_50_ and viability assays, and the cultures were treated with cell lysis buffer to harvest cell lysates.

HNEC ALI cultures were treated with a 2-DG formulation intended for nasal application, consisting of 3.5% 2-DG (213 mM) in citrate buffer (50 mM). Treatments were performed by adding 40 µL of 2-DG or placebo (NaCl) to the apical compartment of each ALI culture. The treatment was either removed after 1 h or left on the cells until sample harvest.

For solution application experiments, HBEC ALI cultures were treated with 3.5% 2-DG (213 mM), 2-FDG (192 mM) or 2-FDM (192 mM) in PBS, or placebo (PBS). The HBEC were treated 1 h before infection and 3 h after infection by adding 40 µL 3.5% 2-DG or placebo to the apical compartment. Subsequent treatments were performed by adding an appropriate volume of 1 M 2-DGA (8.5 µL 2-DG or 7.7 µL 2-FDG/2-FDM) to achieve the same dose per well. This was done to avoid removing any released virus while also preventing diluting the treatment.

Solutions of 2-DG 2-FDG or 2-FDM in PBS were used for both aerosol and solution treatment of Calu-3 and HBEC ALI cultures in experiments involving nebulization. The placebo used was PBS alone. The Vitrocell Cloud α 12 aerosol exposure system paired with Aerogen Pro nebulizers was used for aerosol drug delivery. Aerosol exposure was performed at 34°C, and one treatment consisted of four exposures (1 min nebulization, 10 min sedimentation). For each exposure, 200 µL of solution was nebulized in the 9-well chamber and 76.4 µL in the 3-well chamber. Basal medium cover adapter rings were utilized to ensure that the basal medium was not exposed to nebulized drug. The solution treatment was performed by adding 40 µL 2-DGA or placebo to the apical compartment of each transwell. The concentration of the 2-DGA solution was adjusted to match the expected deposited dose per transwell following nebulization, according to the experimentally determined deposition rate (as previously described ^44^).

### THP-1 activation and NF-κB translocation assay

The THP-1 NF-κB eGFP reporter human monocytic cell line has been genetically modified to express eGFP upon translocation of NF-κB to the nucleus following its activation^33^. The THP-1 cells were seeded into 96-well flat-bottom plates in THP-1 medium at a density of 1.0×10^5^ cells/well. They were pre-treated with 30 mM 2-DG, 2-FDG or 2-FDM or placebo (THP-1 medium) for 1 h, before being activated with 30 µM R848 or treated with vehicle alone (unstimulated control). The 2-DGA treatment was continued after activation for 24 h.

Thereafter, the supernatant was collected for cytokine analysis and the cells were stained with the Zombie Violet Fixable Viability Kit. The cells were washed in PBS and stained in 50 µL viability dye diluted 1:500 in PBS, for 10 min at 4°C. The THP-1 cells were washed and resuspended in PBS before being acquired on a BD LSRFortessa flow cytometer to assess NF-κB translocation. The population of interest was identified by excluding debris, doublets and dead cells, and the data was presented as fold change in eGFP median fluorescence intensity compared to the unstimulated control.

### Cytokine analysis

The LEGENDplex COVID-19 Cytokine Storm Mix & Match kit was used to measure secreted cytokines and chemokines (IL-1β, IL-6, CCL2, CCL5, CXCL8, CXCL10 and TNFα). The analytes were quantified cell culture medium, in technical duplicates, according to the manufacturer’s instructions but adapted to use ¼ volumes.

### RNA isolation, cDNA synthesis and qPCR

Total RNA was extracted from cell lysates using the innuPREP RNA Mini kit 2.0 or QIAGEN RNAeasy Mini Kit, following the manufacturers’ protocol. The RNA concentration of each sample was normalised to 200-400 ng by diluting in nuclease-free water and cDNA was synthesized using the First Strand cDNA Synthesis Kit, using oligo(dT)_18_ primers, according to the manufacturer’s instructions (see Supplementary Table S3 for cycling conditions). qPCR was performed using the PowerTrack SYBR Green Master Mix according to the manufacturer’s instructions (see Supplementary Table S3 for cycling conditions and Supplementary Table S4 for primer sequences). The intracellular viral RNA and *CXCL10* was measured by normalizing the gene expression to host housekeeping gene, hypoxanthine phosphoribosyltransferase 1 (*HPRT*; for HeLa Ohio) or β-2 microglobulin (*B2M;* for ALI cultures) using the 2^-ΔΔCT^ method^45^. The data was expressed as fold changes or percentages, compared to the average of a control group.

### Median tissue culture infectious dose (TCID_50_) assay

For TCID_50_ assays, cells were seeded into 96-well plates in cell-specific infection medium and incubated overnight at 37°C, 5% CO_2_, 85% humidity. HeLa Ohio cells were detached using Accutase and seeded at a density of 1.1-1.5×10^4^ cells per well in 100 µL of HeLa Ohio infection medium. MRC-5 cells were detached with 0.5% trypsin-EDTA and seeded at a density of 1.5-2.0×10^4^ cells per well in 100 µL MRC-5 infection medium. MDCK cells were detached using 1x trypLE and seeded at a density of 1.5-2.0×10^4^ cells per well in 100 µL MDCK infection medium.

Viral supernatants were diluted in full logarithmic steps from 10^-1^ to 10^-11^ or half logarithmic steps from 10^-1^ to 10^-6^ in specific infection medium (as described above). Titrations for RV, HCoV-229E and IAV-H1N1 were conducted in HeLa Ohio, MRC-5 and MDCK cells, respectively. The cells were infected by adding 50 µL of virus-containing supernatants and incubated at 34°C, 5% CO_2_. After 5-6 days the cells were examined under a microscope for cytopathic effect (CPE). For HCoV-229E and IAV-H1N1, CPE or no CPE was documented through microscopic observation. For RV-A16 and RV-A1B, supernatants were aspirated, the wells were washed with PBS, stained with crystal violet solution (0.05% crystal violet powder in 20% methanol/ddH_2_O solution) for 30-60 min and washed three times with ddH_2_O. After drying, 25% acetic acid was added to the wells, the plate was shaken for 2 min and the absorbance at 450 nm was recorded on a plate reader. CPE was defined as an absorbance decrease ≥25% in comparison to the average absorbance of uninfected control wells. The data was analysed according to Reed-Muench’s method^46^.

### Viability assay

The LDH-Glo Cytotoxicity Assay was used according to the manufacturer’s instructions to assess cell viability. Apical wash samples were diluted 3-20X in LDH storage buffer and stored at -20°C until analysis. Maximum LDH release controls were prepared by treating one ALI culture with 40 μL of 10% Triton X-100 for 2 h at 37°C. All samples were normalised to the maximum LDH release control sample, which was considered as 100% cytotoxicity.

### Trans-epithelial electrical resistance (TEER) measurement

To assess barrier integrity, the TEER was measured. The culture medium was aspirated and 200 µL and 500 µL PBS was added to apical and basal compartment, respectively. The cells were incubated at room-temperature for 10 min to equilibrate the temperature. Resistance (R) was measured using an electrode connected to a Millicell-ERS Volt Ohm device. The electrode was held perpendicular and inserted into the transwell without touching the cell layer on the transwell membrane and values were recorded upon stabilisation. The resistance reading from a transwell without cells was used as a blank. The measured resistance values were used to calculate the TEER for each sample (Equation 1):

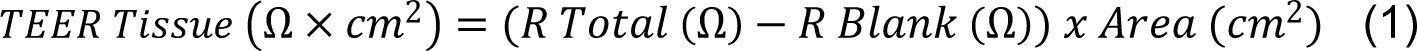

### Cellular 2-FDG uptake assay

Calu-3 cells cultured at ALI were treated with nebulized 3.5% 2-FDG in PBS and the deposited mass was measured using a quartz crystal microbalance. The deposited 2-FDG dose per transwell was calculated by correcting the measured ng/cm^2^ value for the mass stemming from PBS and multiplying by the transwell surface area (0.33 cm2). The calculated value was used to define the solution application dose. Accordingly, Calu-3 cells cultured at ALI were treated with 40 µL of a solution of 2-FDG in PBS, which was added to the apical compartment. The cells were incubated at 34°C, 5% CO_2_, 85% humidity, until metabolite extraction.

At the indicated time points, the solution treatment was removed, and the cells were washed twice with 4°C PBS. Subsequently, 400 µL extraction buffer (50% methanol, 30% acetonitrile, 20% ddH_2_O) was added to the apical compartment and the cells were incubated for 1 hour at 4°C. Finally, the extraction buffer was collected, flash frozen in liquid nitrogen and stored at 80°C until analysis.

To determine the proportion of deposited 2-FDG that was taken up by the cells, a 1-1 conversion rate of 2-FDG to 2-FDG-6P was assumed. The measured amount of 2-FDG-6P in each sample was normalised to the 2-FDG treatment dose as described below (Equations 2 and 3).

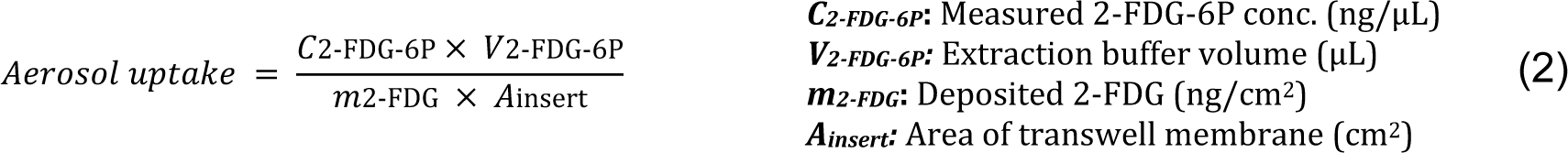

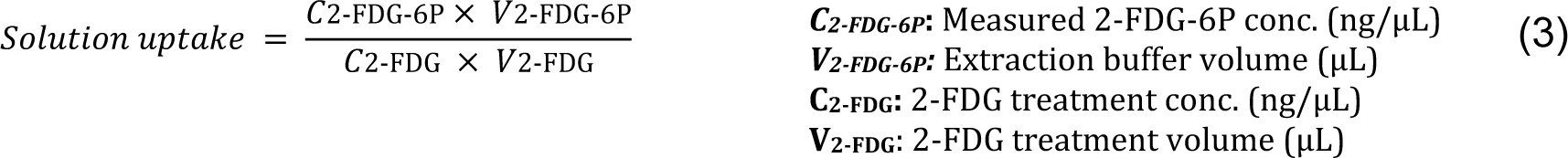

### Mucus penetration assay

Porcine intestine mucus (type III) was used as a substitute for airway mucus, to enable standardization across experiments. PBS was added to each well of the Vitrocell Cloud basal module and empty transwells for aerosol and solution application were placed in separate chambers. Empty transwells without mucus were included in the aerosol group as a control and 20 µL of porcine mucus in PBS (20 mg/mL) was added to the apical compartment of all other transwells. Next, 20 µL of 2-FDG in PBS was added on top of the mucus for the solution application samples. The 2-FDG solution concentration was adjusted to deliver 45 µg per transwell, which is equivalent to the deposited dose following nebulization. Immediately after the solution application, nebulization of 3.5% 2-FDG was initiated for the aerosol application samples. Upon treatment completion, 1000 µL of solution was collected from each basal module well and the samples were stored at -80°C until analysis.

To determine the proportion of 2-FDG that had passed through the mucus layer into the basal compartment, the measured concentration of 2-FDG in each sample was normalised to the highest possible concentration, as described below (Equations 4 and 5):

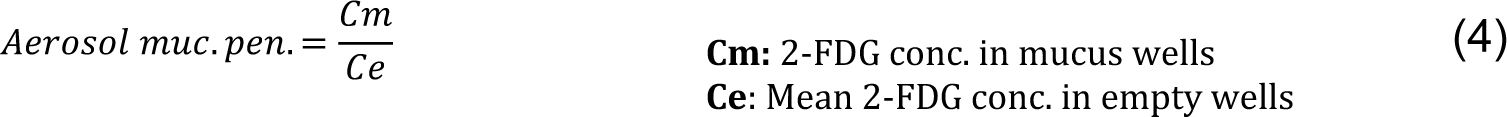

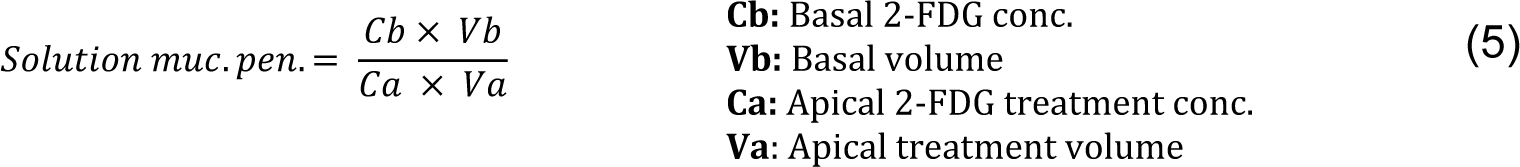

### Quantification of 2-FDG and its metabolite 2-FDG-6P

Samples were analysed by hydrophilic interaction liquid chromatography (HILIC), coupled to tandem mass spectrometry (LC-MS/MS) to measure 2-FDG and 2-FDG-6P.

For 2-FDG measurements, all experimental and calibration curve samples were pretreated by adding 990 µL of extraction solvent (80% acetonitrile, 20% ddH_2_O)) to 10 µL sample volume. The samples were centrifuged and the supernatants further diluted (1:10 - 1:100) to assure a concentration within the linear range of the instrument.

For both 2-FDG and 2-FDG-6P measurements, 1 µL of sample was injected onto a polymeric iHILIC-(P) Classic HPLC column and operated at a flow rate of 100 µL/min (see Supplementary Table S5 for gradients used for separation). An Ultimate 3000 HPLC system was directly coupled via electrospray ionization to a TSQ Quantiva mass spectrometer. Selected reaction monitoring was used for detection and quantification (see Supplementary Table S5 for details). For each transition and metabolite, authentic standards were used to determine optimal collision energies and for the validation of experimental retention times. All samples were measured in technical replicates in a randomized fashion. Injection of pure solvent was used to determine eventual carry over. Data interpretation was performed using TraceFinder (Thermo Fisher Scientific) and absolute values were derived by external calibration.

### Software and statistical analysis

Graphs were generated and statistical analysis was performed using GraphPad Prism (GraphPad Software), flow cytometry data analysis was performed using FlowJo (BD), and experimental setup graphics were created in Inkscape.

Differences were considered statistically different when p<0.05. The Shapiro-Wilk’s normality test was used to determine normality. Parametric statistical tests (e.g. Welch’s t-test) were used for normally distributed data. If the data did not have a normal distribution, or too few data points were available for normality testing, a nonparametric test (e.g. Kruskal-Wallis test) was used. Welch’s correction was applied for t-tests, as it corrects for unequal standard deviations but does not introduce error when standard deviations are equal. An appropriate multiple comparisons correction was used for all statistical tests that compare more than two groups. The applied statistical test, sample size (n) and independent experimental replicates (N) is indicated in each figure legend.

## Data availability statement

All data are available in the main text or in the Supplementary Information.

## Acknowledgements

The authors would like to thank Irene Gössler and Dieter Blaas for their generous provision of viral stocks and Peter Steinberger for kindly providing us with the NF-κB eGFP reporter THP-1 cells. Additionally, we are grateful to G.ST Antivirals employees Sonia Rodriguez, Lena Hell and David Prearo for their advice and support.

## Author contributions

S.K.W., L.W., V.K., S.C., V.G., C.N., H.T. and D.S. performed experiments and analysed data. S.R., A.-D.G., J.S. and G.G. acquired funding, planned and directed the study. S.K.W. and S.R. wrote the manuscript with input from the co-authors. All authors read and approved the final manuscript.

## References

1. Heikkinen, T. & Järvinen, A. The common cold. The Lancet 361, 51–59 (2003).

2. Gern, J. E. How rhinovirus infections cause exacerbations of asthma. Clin. Exp. Allergy 45, 32–42 (2015).

3. Love, M. E. & Proud, D. Respiratory Viral and Bacterial Exacerbations of COPD—The Role of the Airway Epithelium. Cells 11, (2022).

4. Shah, A. et al. Pathogenicity of individual rhinovirus species during exacerbations of cystic fibrosis. Eur. Respir. J. 45, 1745–1748 (2015).

5. Blaas, D. & Fuchs, R. Mechanism of human rhinovirus infections. Mol. Cell. Pediatr. 3, 21 (2016).

6. Liu, D. X., Liang, J. Q. & Fung, T. S. Human Coronavirus-229E, -OC43, -NL63, and -HKU1 (Coronaviridae). in Encyclopedia of Virology 428–440 (Elsevier, 2021). doi:10.1016/B978-0-12-809633-8.21501-X.

7. Keilman, L. J. Seasonal Influenza (Flu). Nurs. Clin. North Am. 54, 227–243 (2019).

8. Gustin, K. M., Belser, J. A., Katz, J. M., Tumpey, T. M. & Maines, T. R. Innovations in modeling influenza virus infections in the laboratory. Trends Microbiol. 20, 275–281 (2012).

9. Miller, E. K. New Human Rhinovirus Species and Their Significance in Asthma Exacerbation and Airway Remodeling. Immunol. Allergy Clin. North Am. 30, 541–552 (2010).

10. Loo, S.-L. et al. Human coronaviruses 229E and OC43 replicate and induce distinct antiviral responses in differentiated primary human bronchial epithelial cells. Am. J. Physiol.-Lung Cell. Mol. Physiol. 319, L926–L931 (2020).

11. Tang, G., Liu, Z. & Chen, D. Human coronaviruses: Origin, host and receptor. J. Clin. Virol. 155, 105246 (2022).

12. Mäkelä, M. J. et al. Viruses and Bacteria in the Etiology of the Common Cold. J. Clin. Microbiol. 36, 539–542 (1998).

13. Matthay, M. A. et al. Acute respiratory distress syndrome. Nat. Rev. Dis. Primer 5, 18 (2019).

14. Traini, D. Inhalation Drug Delivery. Inhalation Drug Delivery: Techniques and Products (Wiley-Blackwell, 2013). doi:10.1002/9781118397145.ch1.

15. Ehrhardt, C. Inhalation Biopharmaceutics: Progress Towards Comprehending the Fate of Inhaled Medicines. Pharmaceutical Research 1–3 (2017) doi:10.1007/s11095-017-2304-2.

16. Fröhlich, E. Biological obstacles for identifying in vitro-in vivo correlations of orally inhaled formulations. Pharmaceutics 11, 1–19 (2019).

17. Radivojev, S., Zellnitz, S., Paudel, A. & Fröhlich, E. Searching for physiologically relevant in vitro dissolution techniques for orally inhaled drugs. Int. J. Pharm. 556, 45–56 (2019).

18. Radivojev, S. et al. Integration of mucus and its impact within in vitro setups for inhaled drugs and formulations: Identifying the limits of simple vs. complex methodologies when studying drug dissolution and permeability. Int. J. Pharm. 661, 124455 (2024).

19. Zellnitz, S. et al. Impact of drug particle shape on permeability and cellular uptake in the lung. Eur. J. Pharm. Sci. 139, 105065 (2019).

20. Fröhlich, E., et al. In vitro toxicity screening of polyglycerol esters of fatty acids as excipients for pulmonary formulations. Toxicol. Appl. Pharmacol. 386, 114833 (2020).

21. Mondoñedo, J. R. et al. A High-Throughput System for Cyclic Stretching of Precision-Cut Lung Slices During Acute Cigarette Smoke Extract Exposure. 11, 1–10 (2020).

22. Pajak, B. et al. 2-Deoxy-d-Glucose and Its Analogs: From Diagnostic to Therapeutic Agents. Int. J. Mol. Sci. 21, 234 (2019).

23. Mayer, K. A., Stöckl, J., Zlabinger, G. J. & Gualdoni, G. A. Hijacking the supplies: Metabolism as a novel facet of virus-host interaction. Front. Immunol. 10, (2019).

24. Wali, L. et al. Host-directed therapy with 2-Deoxy-D-glucose inhibits human rhinoviruses, 1 endemic coronaviruses, and SARS-CoV-2 2 3. J. Virus Erad. (2022) doi:10.1101/2022.05.24.493068.

25. Gualdoni, G. A. et al. Rhinovirus induces an anabolic reprogramming in host cell metabolism essential for viral replication. Proc. Natl. Acad. Sci. 115, (2018).

26. Schmidt, M. F., Schwarz, R. T. & Ludwig, H. Fluorosugars inhibit biological properties of different enveloped viruses. J. Virol. 18, 819–823 (1976).

27. Schlesinger, M. et al. Glucose and mannose analogs inhibit KSHV replication by blocking *N* -glycosylation and inducing the unfolded protein response. J. Med. Virol. 95, e28314 (2023).

28. Kilbourne, E. D. Inhibition of Influenza Virus Multiplication with a Glucose Antimetabolite (2-deoxy-D-glucose). Nature 183, 271–272 (1959).

29. Raez, L. E. et al. A phase I dose-escalation trial of 2-deoxy-d-glucose alone or combined with docetaxel in patients with advanced solid tumors. Cancer Chemother. Pharmacol. 71, 523–530 (2013).

30. Lee, R. E et al. Air-Liquid interface cultures to model drug delivery through the mucociliary epithelial barrier. Adv. Drug Deliv. Rev. 198, 114866 (2023).

31. Lee, D. F., Lethem, M. I. & Lansley, A. B. A comparison of three mucus-secreting airway cell lines (Calu-3, SPOC1 and UNCN3T) for use as biopharmaceutical models of the nose and lung. Eur. J. Pharm. Biopharm. 167, 159–174 (2021).

32. Fröhlich, E. Toxicity of orally inhaled drug formulations at the alveolar barrier: Parameters for initial biological screening. Drug Deliv. 24, 891–905 (2017).

33. Battin, C. et al. A human monocytic NF-κB fluorescent reporter cell line for detection of microbial contaminants in biological samples. PLOS ONE 12, e0178220 (2017).

34. Baldassi, D., Gabold, B. & Merkel, O. M. Air−Liquid Interface Cultures of the Healthy and Diseased Human Respiratory Tract: Promises, Challenges, and Future Directions. Adv. NanoBiomed Res. 1, 2000111 (2021).

35. Ziegler, P. et al. A primary nasopharyngeal three-dimensional air-liquid interface cell culture model of the pseudostratified epithelium reveals differential donor- and cell type-specific susceptibility to Epstein-Barr virus infection. PLOS Pathog. 17, e1009041 (2021).

36. Veerati, P. C. et al. Airway Epithelial Cell Immunity Is Delayed During Rhinovirus Infection in Asthma and COPD. Front. Immunol. 11, 974 (2020).

37. Bhatt, A. N. et al. Glycolytic inhibitor 2-deoxy-d-glucose attenuates SARS-CoV-2 multiplication in host cells and weakens the infective potential of progeny virions. Life Sci. 295, 120411 (2022).

38. Hodes, D. S., Schnitzer, T. J., Kalica, A. R., Camargo, E. & Chanock, R. M. Inhibition of respiratory syncytial, parainfluenza 3 and measles viruses by 2-deoxy-d-glucose. Virology 63, 201–208 (1975).

39. Gualdoni, G. A. et al. Rhinovirus induces an anabolic reprogramming in host cell metabolism essential for viral replication. Proc. Natl. Acad. Sci. U. S. A. 115, E7158– E7165 (2018).

40. O’Neill, L. A. J., Kishton, R. J. & Rathmell, J. A guide to immunometabolism for immunologists. Nat. Rev. Immunol. 16, 553–565 (2016).

41. Lenz, A.-G. et al. Efficient Bioactive Delivery of Aerosolized Drugs to Human Pulmonary Epithelial Cells Cultured in Air–Liquid Interface Conditions. Am. J. Respir. Cell Mol. Biol. 51, 526–535 (2014).

42. Chen, P.-H. et al. Metabolic Diversity in Human Non-Small Cell Lung Cancer Cells. Mol. Cell 76, 838–851.e5 (2019).

43. Cingolani, E. et al. In vitro investigation on the impact of airway mucus on drug dissolution and absorption at the air-epithelium interface in the lungs. Eur. J. Pharm. Biopharm. 141, 210–220 (2019).

44. Bannuscher, A. et al. An inter-laboratory effort to harmonize the cell-delivered in vitro dose of aerosolized materials. NanoImpact 28, 100439 (2022).

45. Livak, K. J. & Schmittgen, T. D. Analysis of relative gene expression data using real-time quantitative PCR and the 2(-Delta Delta C(T)) Method. Methods San Diego Calif 25, 402– 408 (2001).

46. Reed, L. J. & Muench, H. A Simple Method of Estimating Fifty Per Cent Endpoints. Am. J. Hyg. 27, 493–497 (1938).

