## Supplemental Material for "Nebulized 2-deoxylated glucose analogues inhibit respiratory viral infection in advanced *in vitro* airway models"

### Supplementary Information

#### Supplementary Figures

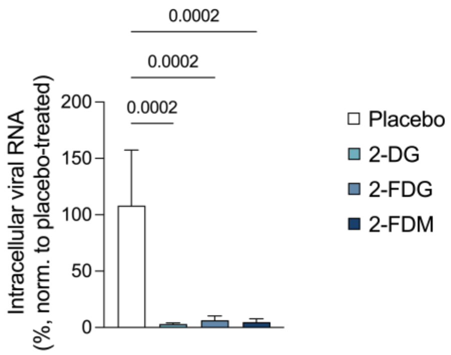


**Supplementary Figure S1**. **2-DGA have antiviral activity in HeLa Ohio cells**. Viral RNA, relative to placebo-treated control, in HeLa Ohio cells infected with RV-B14 and treated with 2-DG, 2-FDG or 2-FDM (10 mM) for 7 hours. Displaying mean, SD, one-way ANOVA with Dunnett's test, n=4, N=2. **Abbreviations:** 2-deoxylated glucose analogues (2-DGA), 2-deoxy-D-glucose (2-DG), 2-fluoro-2-deoxy-D-glucose (2-FDG) and 2-fluoro-2-deoxy-D-mannose (2-FDM), rhinovirus (RV).

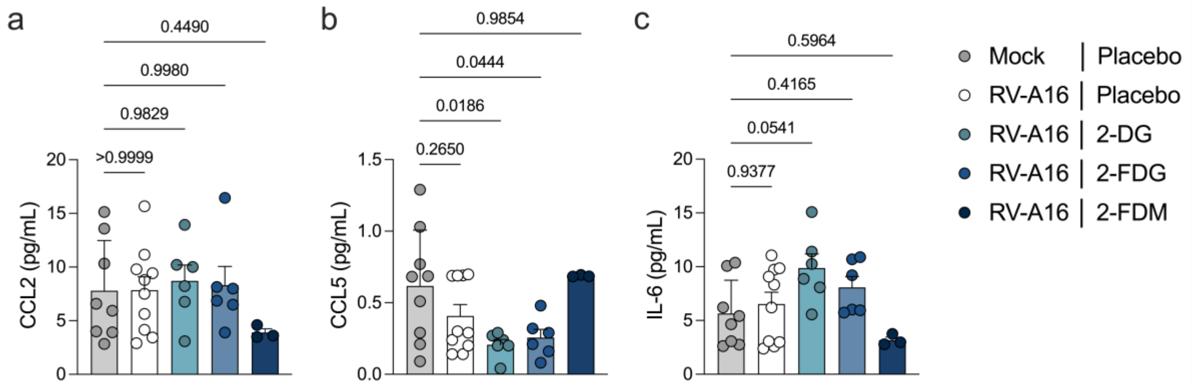


**Supplementary Figure S2. RV infection does not induce CCL2, CCL5 or IL-6 in ALI cultured HBEC.** ALI cultured HBEC were infected with mock or RV-A16 and treated with a solution of 3.5% 2-DG, 2-FDG or 2-FDM at -1, 3, 6, 18, 21 and 24 hours post-infection. (**a**) CCL2, (**b**) CCL5 and (**c**) IL-6 levels in the basal medium 27 hours post-infection. IL-1β and TNFα levels were also measured but were below the detection limit of the assay. **Abbreviations:** 2-deoxy-D-glucose (2-DG), 2-fluoro-2-deoxy-D-glucose (2-FDG), 2-fluoro-2-deoxy-D-mannose (2-FDM), air-liquid-interface (ALI), human bronchial epithelial cells (HBEC), rhinovirus (RV).

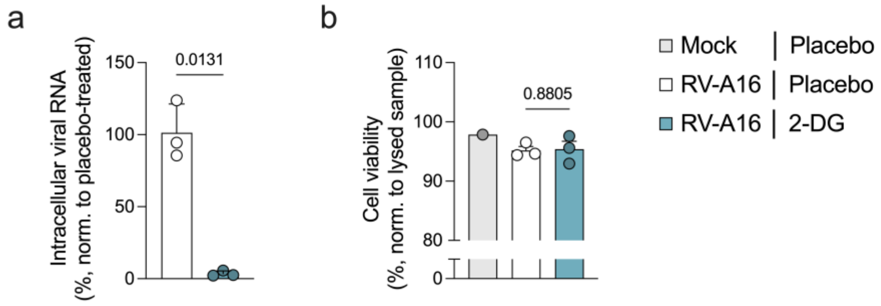


**Supplementary Figure S3. Nebulized 2-DG inhibits RV-A16 replication.** Calu-3 cells cultured at ALI were infected with mock or RV-A16, treated with nebulized 3.5% 2-FDG -1, 2.5, 5 and 7.5 hours post-infection, and samples were harvested 10 hours post-infection. (**a**) RV-A16 viral RNA relative to the placebo-treated control. Displaying mean, SD, Welch's t-test, n=3, N=1. (**b**) Cell viability measured via LDH release and normalised to a lysed Calu-3 sample. Displaying mean, SD, Welch's t-test, n=3, N=1.
**Abbreviations:** 2-deoxy-D-glucose (2-DG), air-liquid-interface (ALI), rhinovirus (RV),

#### Supplementary Methods

**Supplementary table S1**. Cells, viruses and key materials and equipment.

| Item | Supplier | Cat. no. | Additional information |
|---|---|---|---|
| **CELLS** | | | |
| Calu-3 cells (human lung adenocarcinoma origin) | ATCC | HTB-55 | |
| Human nasal epithelial cells (HNEC) | Epithelix Sàrl | EP51AB | Donor no. AB0857 and AB060801 |
| Human bronchial epithelial cells (HBEC) | Epithelix Sàrl | EP51AB | Donor no. 02AB0859 |
| Immortalized human bronchial epithelial cell line (HBEC3-KT) | Evercyte | CkHT-004-0230 | Originally provided by LifeTaq, Austria. |
| HeLa Ohio cells | ECACC | 14J003 | |
| MDCK | ATCC | CCL-34 | |
| MRC-5 cells | ATCC | CCL-171 | |
| THP-1 NF-κB eGFP reporter (human acute monocytic leukemia origin) | n.a. | n.a. | Originally provided by Prof. Peter Steinberger, Medical University of Vienna. |
| **VIRUSES** | | | |

| Item | Supplier | Cat. no. | Additional information |
| --- | --- | --- | --- |
| RV-A16 | ATCC | VR283 | Originally provided by Prof. Dieter Blaas, Max Perutz Labs, University of Vienna. |
| RV-A1B | ATCC | VR1645 | Originally provided by Prof. Dieter Blaas, Max Perutz Labs, University of Vienna. |
| RV-B14 | ATCC | VR284 | Originally provided by Prof. Dieter Blaas, Max Perutz Labs, University of Vienna. |
| HCoV-229E | National Collection of Pathogenic Viruses (UK) | 0310051v | Originally provided by Dr. Nicole Doyle, the Pirbright Institute. |
| IAV-H1N1 | National institute of biological standards and controls (UK) | 19/160 | Strain A/California/7/09 44430 E6. |
| **MATERIALS** | | | |
| PBS (w/o $Ca^{2+}$, $Mg^{2+}$)) | Gibco | 710010-1015 | |
| DPBS (w $Ca^{2+}$, $Mg^{2+}$) | Gibco | 17-516F | |
| Animal component free cell dissociation kit | STEMCELL | 5426 | |
| Accutase | Corning | 25058049 | |
| Trypsin-EDTA | Gibco | R001100 | |
| 2-DG | Sigma | D8375 | |
| 2-FDG | Patheon Regensburg | GDEF-04 | |
| 2-FDM | Patheon Regensburg | GDEF-05 | |
| Type III mucin from porcine stomach | Sigma | M1778 | |
| innuPREP RNA Mini Kit 2.0 | IST Innuscreen GmbH | 845-KS-2040250 | |
| RNeasy Mini Kit | QIAGEN | 74104 | |
| First strand cDNA Synthesis | Thermo Scientific | K1612 | |
| PowerTrack SYBR Green Master Mix | Applied Biosystems | A46109 | |

| Item | Supplier | Cat. no. | Additional information |
|---|---|---|---|
| LDH-Glo Cytotoxicity Assay | Promega | J2380 / J2381 | |
| LEGENDplex COVID-19 Cytokine Storm Mix & Match | BioLegend | 741089 / 741112 | |
| Zombie Violet Fixable Viability Kit | BioLegend | 423113 | |
| Costar® 6.5 mm Transwell®, 0.4 µm Pore Polyester Membrane Inserts | Corning | 3470 | |
| iHILIC-(P) Classic HPLC column (200x2.1 mm; 5 µm) | HILICON | n.a. | Used for 2-FDG measurements. |
| iHILIC-(P) Classic HPLC column (100x2.1 mm; 5 µm) | HILICON | n.a. | Used for 2-FDG-6P measurements. |
| **EQUIPMENT** | | | |
| Cloud $\alpha$ 12 exposure system | Vitrocell Systems | n.a. | |
| sQCM 12 Quartz Crystal Microbalance | Vitrocell Systems | n.a. | |
| Aerogen Pro nebulizer | Aerogen | 90-002 | |
| Millicell-ERS Volt-Ohm meter | Merck | MERS00002 | |
| QuantStudio 1 Real-Time PCR Instrument | Applied Biosystems | A40425 | |
| Nanophotometer N50-Touch | IMPLEN | T51384 | |
| GENios / Spark microplate reader | TECAN | n.a. | |
| Ultimate 3000 HPLC system; | Dionex, Thermo Fisher Scientific | n.a. | |
| TSQ Quantiva mass spectrometer | Thermo Fisher Scientific | n.a. | |

**Supplementary Table S2**. Cell culture medium composition.

| Medium name | Component | Concentration | Supplier | Cat. No. |
|---|---|---|---|---|
| HeLa Ohio maintenance medium | RPMI 1640 medium (2 g/L glucose) | 1X | Gibco | 21875034 |
| | FBS | 10% | Gibco | 10270106 |
| | penicillin/streptomycin | 100 Units/mL | Gibco | 15140122 |
| | L-glutamine | 2 mM | Gibco | 25030024 |
| | MycoZAP | 0.2% | Lonza | VZA-2032 |
| HeLa Ohio experiment medium | RPMI 1640 medium (0 g/L glucose) | 0.5X | Gibco | 11879020 |
| | RPMI 1640 medium (2 g/L glucose) | 0.5X | Gibco | 21875034 |
| | FBS | 10% | Gibco | 10270106 |
| | Penicillin/streptomycin | 100 Units/mL | Gibco | 15140122 |
| | L-glutamine | 2 mM | Gibco | 25030024 |
| HeLa Ohio infection medium | MEM (1 g/L glucose) | 1X | Gibco | 11514426 |
| | FBS | 2% | Gibco | 10270106 |
| | Penicillin/streptomycin | 100 Units/mL | Gibco | 15140122 |
| | L-glutamine | 2 mM | Gibco | 25030024 |
| | $MgCl_3$ | 30 mM | Thermo Fisher Scientific | 223211000 |
| MRC-5 maintenance medium | MEM (1 g/L glucose) | 1X | Gibco | 11514426 |
| | FBS | 10% | Gibco | 10270106 |
| | Penicillin/streptomycin | 100 Units/mL | Gibco | 15140122 |
| | Sodium pyruvate | 1 mM | Gibco | 11360-070 |
| MRC-5 infection medium | MEM (1 g/L glucose) | 1X | Gibco | 11514426 |
| | FBS | 2% | Gibco | 10270106 |
| | Penicillin/streptomycin | 100 Units/mL | Gibco | 15140122 |
| | Sodium pyruvate | 1 mM | Gibco | 11360-070 |

| Medium name | Component | Concentration | Supplier | Cat. No. |
|---|---|---|---|---|
| MDCK maintenance medium | MEM (1 g/L glucose) | 1X | Gibco | 11514426 |
| | FBS | 10% | Gibco | 10270106 |
| | Sodium pyruvate | 1 mM | Gibco | 11360-070 |
| MDCK infection medium | MEM (1 g/L glucose) | 1X | Gibco | 11514426 |
| | Sodium pyruvate | 1 mM | Gibco | 11360-070 |
| | TrypLE | 1% | Gibco | 12563 |
| THP-1 medium | RPMI 1640 medium (2 g/L glucose) | 1X | Gibco | 21875034 |
| | FBS (heat-inactivated for 10 min at 56 °C) | 10% | Gibco | 10270106 |
| | Penicillin/streptomycin | 100 Units/mL | Gibco | 15140122 |
| PneumaCult-Ex Plus medium | PneumaCult-Ex Plus basal medium | 1X | STEMCELL | 05040 |
| | PneumaCult-Ex Plus 50X supplement | 1X | STEMCELL | 05040 |
| | Hydrocortisone stock solution | 0.2 µM | STEMCELL | 07925 |
| | Penicillin/streptomycin | 100 Units/mL | Gibco | 15140122 |
| PneumaCult-ALI medium | PneumaCult-ALI basal medium | 1X | | 05001 |
| | PneumaCult-ALI 10X supplement | 1X | | 05001 |
| | PneumaCult-ALI 100X supplement | 1X | | 05001 |
| | Hydrocortisone stock solution | 1 µM | STEMCELL | 07925 |
| | Penicillin/streptomycin | 100 Units/mL | Gibco | 15140122 |
| | Heparin | 0.72 Units/mL | STEMCELL | 07980 |

| Medium name | Component | Concentration | Supplier | Cat. No. |
|---|---|---|---|---|
| HBEC infection medium | RPMI 1640 medium (0 g/L glucose) | 0.5X | Gibco | 11879020 |
| | PneumaCult-ALI basal medium | 0.5X | | 05001 |
| | PneumaCult-ALI 10X supplement | 1X | | 05001 |
| | PneumaCult-ALI 100X supplement | 1X | | 05001 |
| | Hydrocortisone stock solution | 1 µM | STEMCELL | 07925 |
| | Penicillin/streptomycin | 100 Units/mL | Gibco | 15140122 |
| | L-glutamine | 1 mM | Gibco | 25030024 |
| | Heparin | 0.72 Units/mL | STEMCELL | 07980 |
| Keratinocyte-SFM medium | Keratinocyte-SFM basal medium | 1X | Gibco | 17005-042 |
| | Bovine pituitary extract | 0.05 mg/mL | Gibco | 17005-042 |
| | Human recombinant epidermal growth factor | 0.005µg/mL | Gibco | 17005-042 |
| | Penicillin/streptomycin | 100 Units/mL | Gibco | 15140122 |
| Calu-3 medium | MEM (1 g/L glucose) | 1X | Gibco | 11514426 |
| | FBS | 10% | Gibco | 10270106 |
| | Penicillin/streptomycin | 100 Units/mL | Gibco | 15140122 |
| | L-glutamine | 2 mM | Gibco | 25030024 |

**Supplementary Table S3.** Cycling times and temperatures.

| Application | Step | Temperature (°C) | Time (min) | Details |
|---|---|---|---|---|
| cDNA synthesis | 1 | 65 | 5 | Oligo(dT)$_{18}$ primer and RNA only. |
| | 2 | 4 | 5 | |
| | 3 | 37 | 60 | After the addition of master mix: reaction buffer, RiboLock RNase inhibitor, dNTPs and M-MuLV reverse transcriptase. |
| | 4 | 70 | 5 | |
| RT-qPCR | 1 | 95 | 5 | |
| | 2 | 95 | 0.25 | Repeated for a total of 39 cycles. |
| | 3 | 60 | 0.25 | |
| | 4 | 72 | 0.75 | |
| | 5 | 95 | 0.25 | |
| | 6 | 60 | 1 | |
| | 7 | 95 | 0.25 | |

**Supplementary Table S4.** Primer sequences.

| Name | **Sequence** (5'-3') | Supplier |
|---|---|---|
| B2M_F | TGACTTTGTCACAGCCCAAG | Microsynth |
| B2M_R | TCCAATCCAAATGCGGCATC | Microsynth |
| HPRT_F | TCAGGCAGTATAATCCAAAGATG | Microsynth |
| HPRT_R | AGTCTGGCTTATATCCAACACTT | Microsynth |
| RVB14_F | GGCGCCATATCCAATGGTGT | Microsynth |
| RVB14_R | TCCACCTGATCGAACGTCCA | Microsynth |
| RV-A16_F | CCAAACACACCCAATACTGCTG | Microsynth |
| RV-A16_R | TTGGTCCAGTTTGCTTGTGC | Microsynth |
| CXCL10_F | ACTGCCATTCTGATTTGCTGCCTT | Microsynth |
| CXCL10_R | ACTAATGCTGATGCAGGTACAGCG | Microsynth |

**Supplementary Table S5**. Methodological details for LC-MS.

| HPLC GRADIENTS | Solvent | Transition |
|---|---|---|
| 2-FDG | A: 100% acetonitrile | A linear gradient starting with 22% B and ramping up to 80% B in 18 min was used for separation. |
| | B: 10 mM aqueous ammonium acetate | |
| 2-FDG-6P | A: 100% acetonitrile | A linear gradient starting with 13% B and ramping up to 80% B in 14 minutes was used for separation. |
| | B:10 mM aqueous ammonium bicarbonate, with 0.1 µg/ml medronic acid | |
| **SRM TRANSITIONS** | **Compound measured** | **Ion mass to charge (m/z)** |
| 2-FDG | Acetate adduct of glucose | 239-179 |
| | Acetate adduct of 2-DG | 223-85; 145 |
| | Acetate adduct of 2-FDG and 2-FDM | 241-103 |
| | Qualifier of 2-DG | 163-85; 145 |
| | Qualifier of 2-FDG and 2-FDM | 181-103 |
| 2-FDG-6P | Taurine | 124-80 |
| | Glucose-6-phosphate | 259-97 |
| | Deoxyglucose-6-phosphate | 243-97 |
| | Fluorodeoxyhexose-6-phosphate | 261-97 |
